# Annotation and differential analysis of alternative splicing using *de novo* assembly of RNAseq data

**DOI:** 10.1101/074807

**Authors:** Clara Benoit-Pilven, Camille Marchet, Emilie Chautard, Leandro Lima, Marie-Pierre Lambert, Gustavo Sacomoto, Amandine Rey, Cyril Bourgeois, Didier Auboeuf, Vincent Lacroix

**Affiliations:** Université de Lyon, ENS de Lyon, Université Claude Bernard, CNRS UMR 5239, INSERM U1210, Laboratory of Biology and Modelling of the Cell, 46 Allée d’Italie Site Jacques Monod, F-69007, Lyon, France; Université de Lyon, F-69000, Lyon; Université Lyon 1; CNRS, UMR5558, Laboratoire de Biométrie et Biologie Evolutive, F-69622, Villeurbanne, France. EPI ERABLE - Inria Grenoble - Rhône-Alpes; IRISA Inria Rennes Bretagne Atlantique CNRS UMR 6074, Université Rennes 1, GenScale team, Rennes, 263 Avenue Général Leclerc, Rennes, France

**Keywords:** “*de novo assembly*”, “alternative splicing”, “transcriptome”

## Abstract

Genome-wide analyses reveal that more than 90% of multi exonic human genes produce at least two transcripts through alternative splicing (AS). Various bioinformatics methods are available to analyze AS from RNAseq data. Most methods start by mapping the reads to an annotated reference genome, but some start by a *de novo* assembly of the reads. In this paper, we present a systematic comparison of a mapping-first approach (FaRLine) and an assembly-first approach (KisSplice). These two approaches are event-based, as they focus on the regions of the transcripts that vary in their exon content. We applied these methods to an RNAseq dataset from a neuroblastoma SK-N-SH cell line (ENCODE) differentiated or not using retinoic acid. We found that the predictions of the two pipelines overlapped (70% of exon skipping events were common), but with noticeable differences. The assembly-first approach allowed to find more novel variants, including novel unannotated exons and splice sites. It also predicted AS in families of paralog genes. The mapping-first approach allowed to find more lowly expressed splicing variants, and was better in predicting exons overlapping repeated elements. This work demonstrates that annotating AS with a single approach leads to missing a large number of candidates. We further show that these candidates cannot be neglected, since many of them are differentially regulated across conditions, and can be validated experimentally. We therefore advocate for the combine use of both mapping-first and assembly-first approaches for the annotation and differential analysis of AS from RNAseq data.

## 1 Introduction

In the last 10 years, the prevalence of alternative splicing has been completely re-evaluated. Recent reports claim that more than 90% of multi-exon genes produce at least two splicing variants [Pan et al., 2008, Wang et al., 2008]. The depth at which we can sample transcriptomes with next generation sequencing techniques opens the possibility not only to annotate splicing variants in physiological conditions, but also to detect which transcripts are differentially spliced across conditions.

This growing interest in splicing both as a fundamental process and because of its implication in pathologies [Scotti and Swanson, 2016, Edery et al., 2011, David and Manley, 2010] has been accompanied by an increasing number of methods aiming at analyzing RNAseq datasets [Trapnell et al., 2012, Wang et al., 2010, Robertson et al., 2010]. The ultimate goal of these methods is to identify and quantify full-length transcripts from short sequencing reads. This task is particularly challenging and recent benchmarks show that all methods still make a lot of mistakes [Steijger et al., 2013]. The difficulty of reconstructing full-length transcripts (isoform-centric approaches) also prompted a number of authors to focus on identifying exons that are differentially included within transcripts (exon-centric approaches) [Reyes et al., 2013, Katz et al., 2010, Shen et al., 2012, Sacomoto et al., 2012].

Whether they are exon-centric or isoform-centric, methods to study splicing from RNAseq data can further be divided in two main categories [Martin and Wang, 2011]. The mapping-first approaches first map the reads to the reference genome and the mapped reads are then assembled into exons and eventually transcripts. In contrast, assembly-first approaches first assemble the reads based on their overlaps. The assembled sequences (corresponding to sets of exons) are then aligned to the reference genome.

Mapping-first approaches have been the most used so far, essentially because they were the first to be developed and because they initially required less computational resources. *De novo* assembly methods were also thought to be restricted to non-model species, where no (good) reference genome is available, and they seemed to be inadequate when an annotated reference genome is available.

Recent progress in *de novo* transcriptome assembly is clearly changing this view, and the argument of the heavier computational burden does not hold anymore.

The application of *de novo* assembly to human RNAseq data however still remains rare, although some studies have already shown its potential to detect novel splicing variants which play a central role in the studied disease [Dargahi et al., 2014, Freyermuth et al., 2016].

The generalization of *de novo* assembly approaches for studying splicing in human seems to be mostly impeded by the lack of a clear evaluation of its potential in comparison to more traditional mapping-based approaches.

This is the gap we aim at filling with the work presented here.

To achieve this goal, we performed a systematic evaluation of an assembly-first and a mapping-first approach on the same publicly available RNAseq dataset.

As a first step, we chose to compare pipelines that we developed in parallel in two teams, namely KisSplice and FaRLine, because we could easily control their parameters. Any difference between the predictions that is solely due to a parameter setting could be fixed easily, which enabled us to obtain a precise understanding of the irreducible differences between the two approaches.

In a second step, we benchmarked our methods to other classically used pipelines and were able to confirm the generality of our findings.

A significant part of our work has been to manually dissect a number of cases found by only one of the two methods. This enabled us to go beyond a simple qualitative description and provide the community with a precise understanding of which cases are overlooked by each type of method, and where new methods are needed.

From a general point of view, the combination of approaches we propose will enable researchers to extend significantly their list of candidates.

## 2 Results

### 2.1 KisSplice and FaRLine

Figure 1 presents schematically the two pipelines that we developed and compared. A detailed description of each step is given in the Methods section. In the assembly-first approach, a De Bruijn graph is built from the reads. Alternative splicing events, which correspond to bubbles in this graph are enumerated and quantified by KisSplice. Each path is then mapped on the reference genome using STAR and the event is annotated by KisSplice2RefGenome using EnsEMBL r75 annotations. In the mapping-first approach, reads are aligned to the reference genome using TopHat2. Mapped reads are then analyzed by FaRLine, in the light of the EnsEMBL r75 annotations.

**Figure 1:**
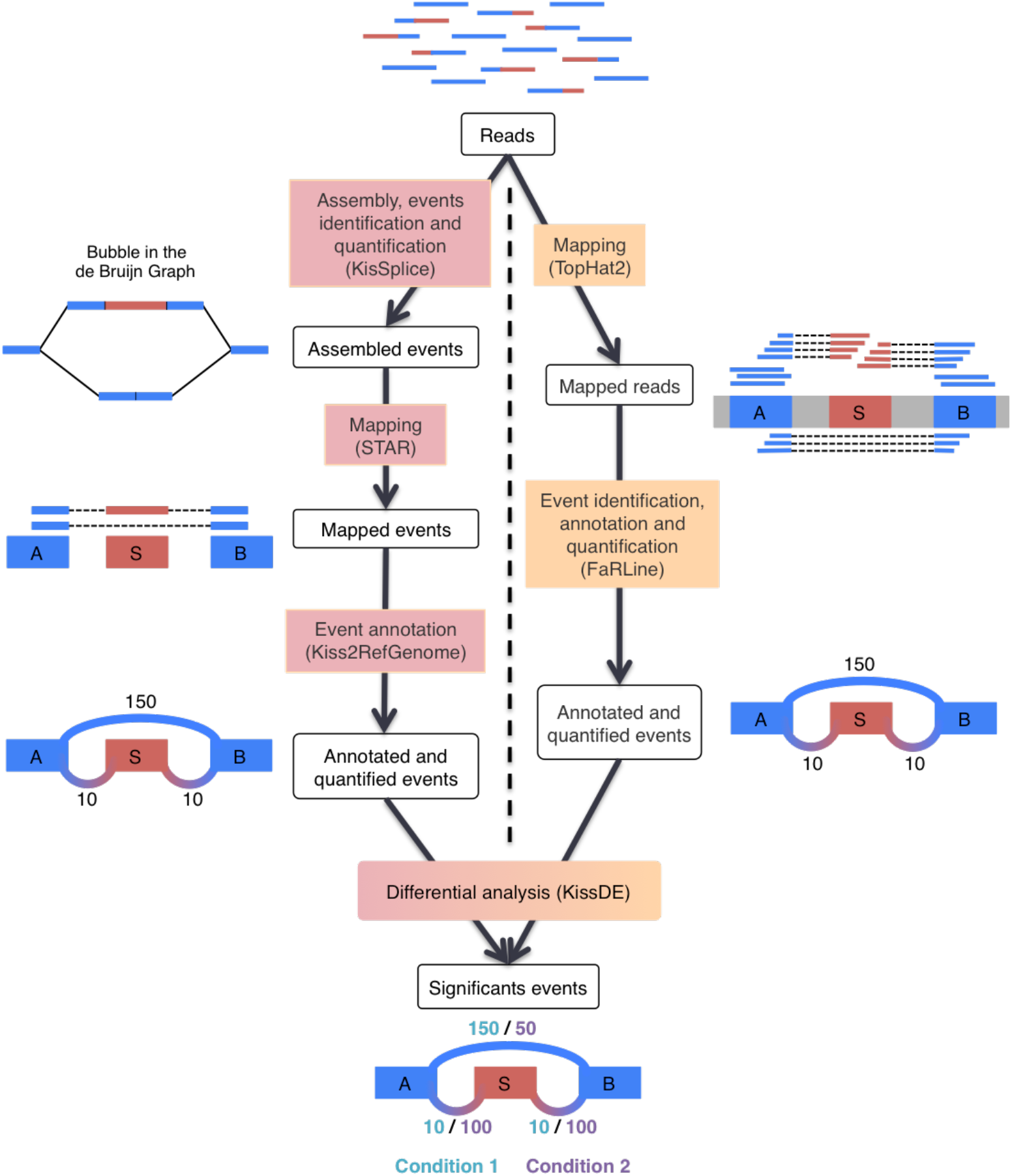
The two pipelines compared in this study: KisSplice and FaRLine. The first step of KisSplice is to assemble the reads and extract the splicing events. These events are then mapped back to the reference genome and classified by event type. The annotated and quantified events are then used for the differential analysis between the biological conditions. In contrast, the first step of FaRLine is to map the reads on the reference genome. From this mapping, annotated and quantified events are extracted. Finally, the differential analysis is done with the same method as in the KisSplice pipeline.

We also tested STAR instead of TopHat2 for the mapping-first pipeline, and found that our main results were essentially unchanged (see Methods).

Quantification of splicing variation is performed similarly in the two pipelines where only junction reads are considered. For the inclusion isoform, there are two junctions to consider. We calculate the mean of the counts of these two junctions.

The differential analysis is performed by a common method for the two approaches: kissDE, which tests if the relative abundance of the inclusion isoform has changed significantly across conditions.

Overall, we further developed and adapted jointly these two pipelines in order to minimize the discrepancies that could unnecessarily complicate our comparison.

### 2.2 The majority of frequent isoforms are found by both approaches

Applying KisSplice and FaRLine to the same RNAseq dataset (SK-N-SH cell lines treated or not with retinoic acid) generated by the ENCODE consortium, we noticed that 68% of the alternatively skipped exons (ASE) identified by KisSplice were also identified by FaRLine and that 24% of ASEs identified by FaRLine were also identified by KisSplice (Figure 2 A). This observation highlights that the mapping-first approach predicts a much larger number of events. This difference in sensitivity is due to the fact that while mapping-first approaches require that each exon junction is covered by at least one read, assembly-first approaches require overlapping reads across the full skipped exon. Therefore, it can be anticipated that low abundant isoforms, that are covered by few reads, will be reported by mapping, but not by the assembly-first approach. Supporting this prediction, we found that for ASEs reported only by FaRLine, the number of reads supporting the minor isoform is much lower than in the other categories (Figure 2 B).

**Figure 2:**
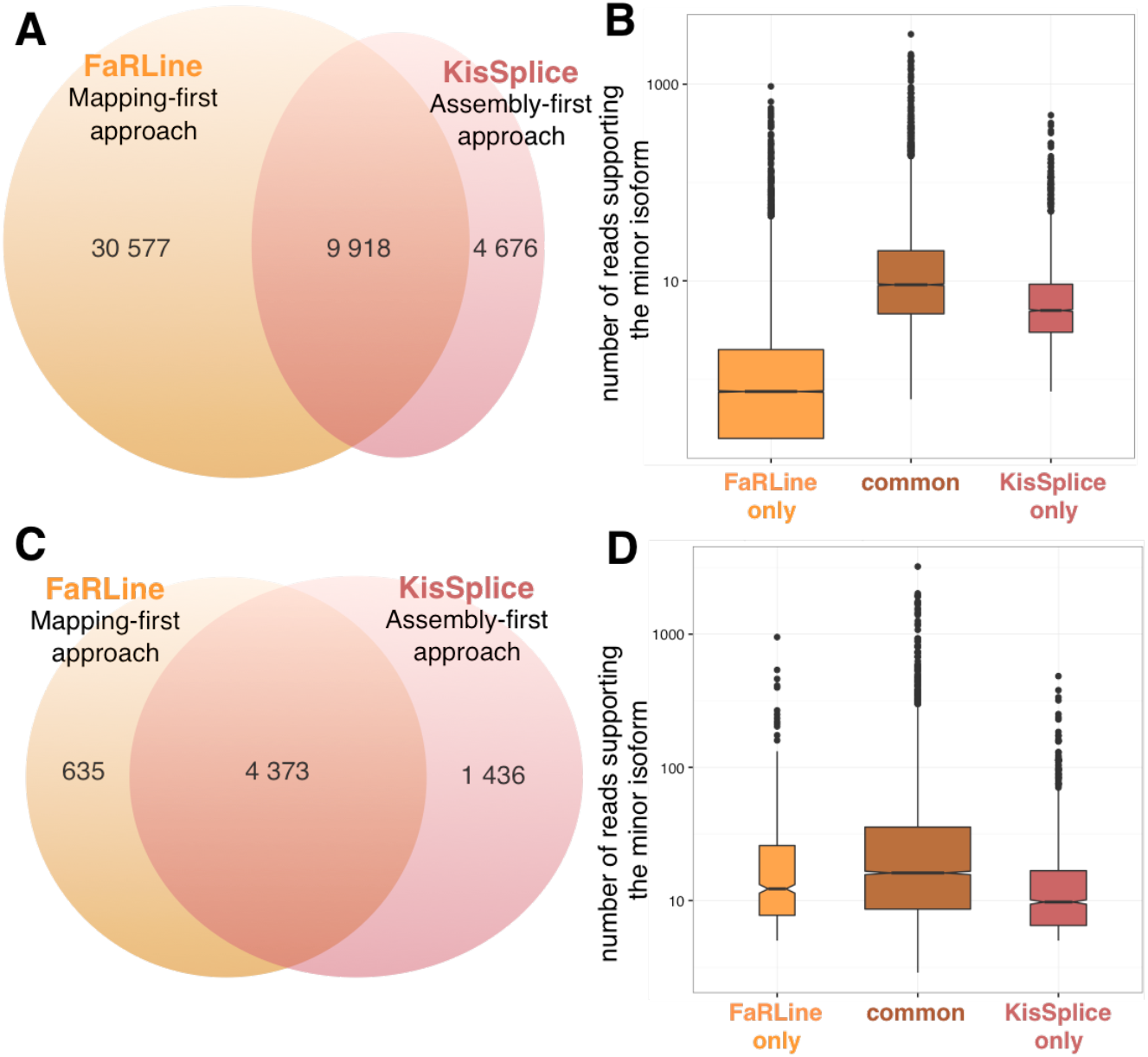
Comparison of the annotated ASE between assembly-first and mapping-first pipelines. A) Venn diagram of ASEs annotated by the two pipelines. FaRLine detected many more events than KisSplice. 68% of ASE annotated by KisSplice were also found by FaRLine and 24% of ASE annotated by FaRLine were also found by KisSplice. B) Boxplot of the expression of the minor isoform in the 3 categories defined in the Venn diagram of panel A: ASE found only by FaRLine, ASE found by both pipelines and ASE found only by KisSplice. The number of reads supporting the minor isoform of the ASE found by FaRLine is globally much lower. C) Venn diagram of ASEs annotated by the two pipelines after filtering out the poorly expressed isoforms. The common events represent a larger proportion of the annotated events than previously: 87% of the ASE annotated by FaRLine and 75% of the ASE annotated by KisSplice. D) Boxplot of the expression of the minor isoform in the 3 categories defined in the Venn diagram of panel C: ASE found only by FaRLine, ASE found by both pipelines and ASE found only by KisSplice. The distribution of the number of reads supporting the minor isoform is similar for the 3 categories with highly expressed variants in each category.

In order to further compare the mapping and assembly-first approaches, we decided to filter out candidates for which the minor isoform was supported by less than 5 reads or whose relative abundance was lower than 10% compared to the major isoform.

As expected, the proportion of candidates reported by both methods increased significantly. Approximately 70% of predicted skipped exons were now found by both approaches. (Figure 2C).

Furthermore, the estimation of their inclusion levels were very consistent across the two approaches (*R*^2^ > 0.9).

Beyond the overall concordance of the two approaches in detecting common splicing events, a number of candidates remained reported by only one approach. Since many of them have a highly-expressed minor isoform (supported by more than 100 reads) (Figure 2D), the failure of one approach to detect them is likely not due to a lack of coverage.

Moreover, events from each of these 3 categories were validated by RT-PCR (Figure 3 and Supplementary Figure S1).

**Figure 3:**
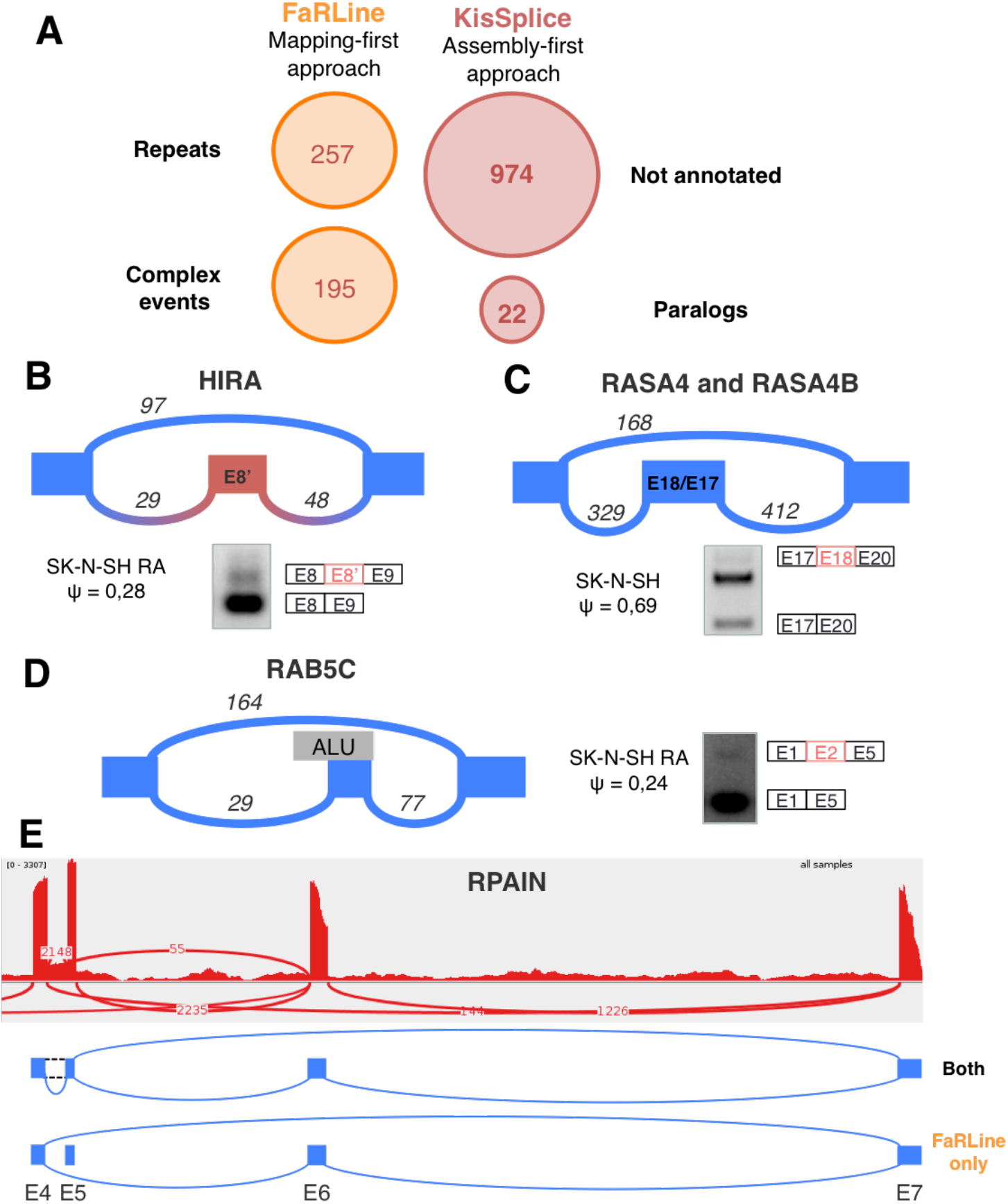
A) Categories identified explaining why some exons are detected by only one method. B) The new exon in intron 8 of the gene HIRA is an example of an exon not annotated in EnsEMBL r75. This event was found by KisSplice but not by FaRLine. C) RASA4 and RASA4B are 2 paralog genes. KisSplice detected 2 isoforms that could be produced by these 2 genes. FaRLine did not find any event in either of these genes. The exon skipped is exon 18 in RASA4 (corresponding to exon 17 in RASA4B). The third band on the RT-PCR is the inclusion of another exon in the intron 18 of RASA4. C) Exon 2 of the gene RAB5C is an example of exon skipping overlapping an Alu found only by FaRLine. The events in panel A to C were validated by RT-PCR. E) RPAIN contains a complex event with a lowly expressed isoform. This weakly expressed isoform was not found by KisSplice, while the other isoforms were found by both approaches.

For all these cases, we patiently dissected the reasons why they could have been missed out by one approach. This enabled us to define 4 main categories (Figure 3A).

### 2.3 Some isoforms are systematically missed by one approach

The first category corresponds to cases that were missed out by the mapping-first approach and corresponds to alternative splicing events involving novel unannotated exons. The unannotated exon can be the skipped exon or one of its flanking exons. It can also be a subpart of a larger annotated exon, and hence be overseen (see Methods).

The reason why the mapping-first approach does not detect these events is twofold. First, the mapper may map the reads to an incorrect location, as junction discovery using short reads is a challenging task. This occurred in 17% of the 1436 cases. Second, in the case where the mapper succeeds (83% of the cases), FaRLine failed to report the event because it relies on annotations. Among these 1199 cases, we distinguished 3 sub-categories of errors due to the annotation. Either the exon is unannotated (28%), one of its flanking exon is unannotated (8%) or both exons are annotated but no transcript combining them was annotated (45%). The assembly-first approach, KisSplice, does not consider annotations, and an interesting resulting advantage is that novel junctions have the same chance to be assembled as known junctions. Mapping assembled novel junctions to the genome is indeed less challenging than read mapping because the assembled sequences are longer.

The downstream annotation of the events is then permissive, in the sense that annotations are used as an evidence, not as a guide. Alternative splicing events involving novel splice sites are clearly identified as such, and can be individually tested and experimentally validated. HIRA gene contain a novel exon, whose inclusion is supported by at least 20 reads on each junction (Figure 3B). This case was overseen by the mapping-first approach, FaRLine. The panel A of the supplementary figure S2 shows an example of an ASE not reported by FaRLine because the inclusion was not present in the transcripts.

The second category of splicing events identified by only one approach corresponds to paralog genes. Untangling the relation between alternative splicing and gene duplication is a difficult topic, subject to debate [Kopelman et al., 2005, Roux and Robinson-Rechavi, 2011]. It is indeed difficult to assess the amount of alternative splicing that occurs within paralogous genes. With the mapping-first approach, the reads stemming from recent paralogs are classified as multi-mapping reads. FaRLine, like the vast majority of mapping-first pipelines, discards these reads for further analysis, as their precise location cannot be clearly established. This results in silently underestimating alternative splicing in paralog genes. In opposition, *de novo* assembly can faithfully state that a family of recent paralogs collectively produce two isoforms that vary in their sequence. However, whether the two isoforms are produced from the same locus or from different loci remains undetermined. KisSplice detects these cases of putative AS in paralog genes. Figure 3C illustrates the case with genes RASA4 and RASA4B. Exon 18 in RASA4 (denoted as exon 17 in RASA4B) was detected to be skipped. The exclusion isoform is supported by 160 reads, while the inclusion isoform is supported by 400 reads. The mapping-first approach did not detect either of these isoforms at all.

The third category of splicing events identified by only one approach corresponds to cases that are missed out by the assembly-first approach. Out of the 635 cases belonging to this category, a large fraction (40%) corresponds to cases where the skipped exon overlaps a repeat, notably Alu elements. Alu are transposable elements present in a very large number of copies in the human genome [Batzer and Deininger, 2002]. Most of these copies are located in introns and a number of them have been exonised [Lev-Maor et al., 2003, Sorek et al., 2004]. The reason why the mapping-first approach is able to identify these cases is because even though the read partially map to repeated sequences, the boundaries of these exons are unique and annotated. Hence the mapper, if set properly, can map these reads to unique annotated exon junctions and is not confused by multiple mappings. Importantly, if the annotations are not provided to the mapper, it will be confused by multiple mappings and will not be able to map the read to the correct location (Supplementary Figure S3). The assembly-based approach fails to detect most of these events. The reason is that, although they do form bubbles in the DBG generated by the reads, these bubbles are highly branching (online supplementary figure http://kissplice.prabi.fr/sknsh/graph_RAB5C_distance_3.html). Enumerating branching bubbles is computationally very challenging, and may take a prohibitive amount of time. In practice, we restrict our search to the enumeration of bubbles with at most 5 branches (Supplementary Figure S4). Increasing this threshold would lead to an increase in the sensitivity at the expense of the running time.

The fourth category of splicing events identified by only one approach corresponds to the cases where more than two splicing isoforms locally coexist, and one of them is poorly expressed compared to the others. The RPAIN gene is a good illustration of such cases (Figure 3E), as exons 5 and 6 of RPAIN may be skipped and the intron between exons 4 and 5 may be retained. While both methods successfully reported the skipping of exon 6, with exons 5 and 7 as flanking, FaRLine additionally reported the skipping of the same exon, but with exons 4 and 7 as flanking exons. The reason why KisSplice did not report this case is because the junction between exons 4 and 6 is relatively weakly supported. More specifically, this junction is supported by only 55 reads, which accounts for less than 2% of the total number of reads branching out from exon 4. Transcriptome assemblers, like KisSplice, usually interpret such relatively weakly supported junctions as sequencing errors or spurious junctions in highly-expressed genes, therefore disregarding them in the assembly phase (see Methods).

### 2.4 Comparison of the approaches after differential analysis

Beyond the tasks of identifying exon skipping events, a natural question which arises when two conditions are compared is to assess if the inclusion level of the exon significantly changed across conditions.

In order to test this, we took advantage of the availability of replicates for both the SK-N-SH cell line and the same cell line treated retinoic acid. For each detected event, we tested with kissDE [Lopez-Maestre et al., 2016], whether we could detect a significant association between one isoform and one condition. Focusing on those condition-specific events, we again partitioned them in events reported by both methods, by FaRLine only and by KisSplice only. As shown in Figure 4, we found again that the majority of condition-specific events were detected by both approaches. This is the case for instance of exon 22 of gene ADD3 which is clearly more included upon retinoic acid treatment (Figure 4C), with a DeltaPSI of 27%. The estimation of the DeltaPSI is overall very similar across the two approaches (Figure 4B) with a correlation of 0.94. The outliers essentially correspond to ASE with several alternative donor/acceptor sites. KisSplice considers these events as different exons while FaRLine considers them as an unique exon, and sums up all the incoming (resp. outgoing) junction counts. Hence, the read counts will differ. Supplementary Figure S5 gives an example.

**Figure 4:**
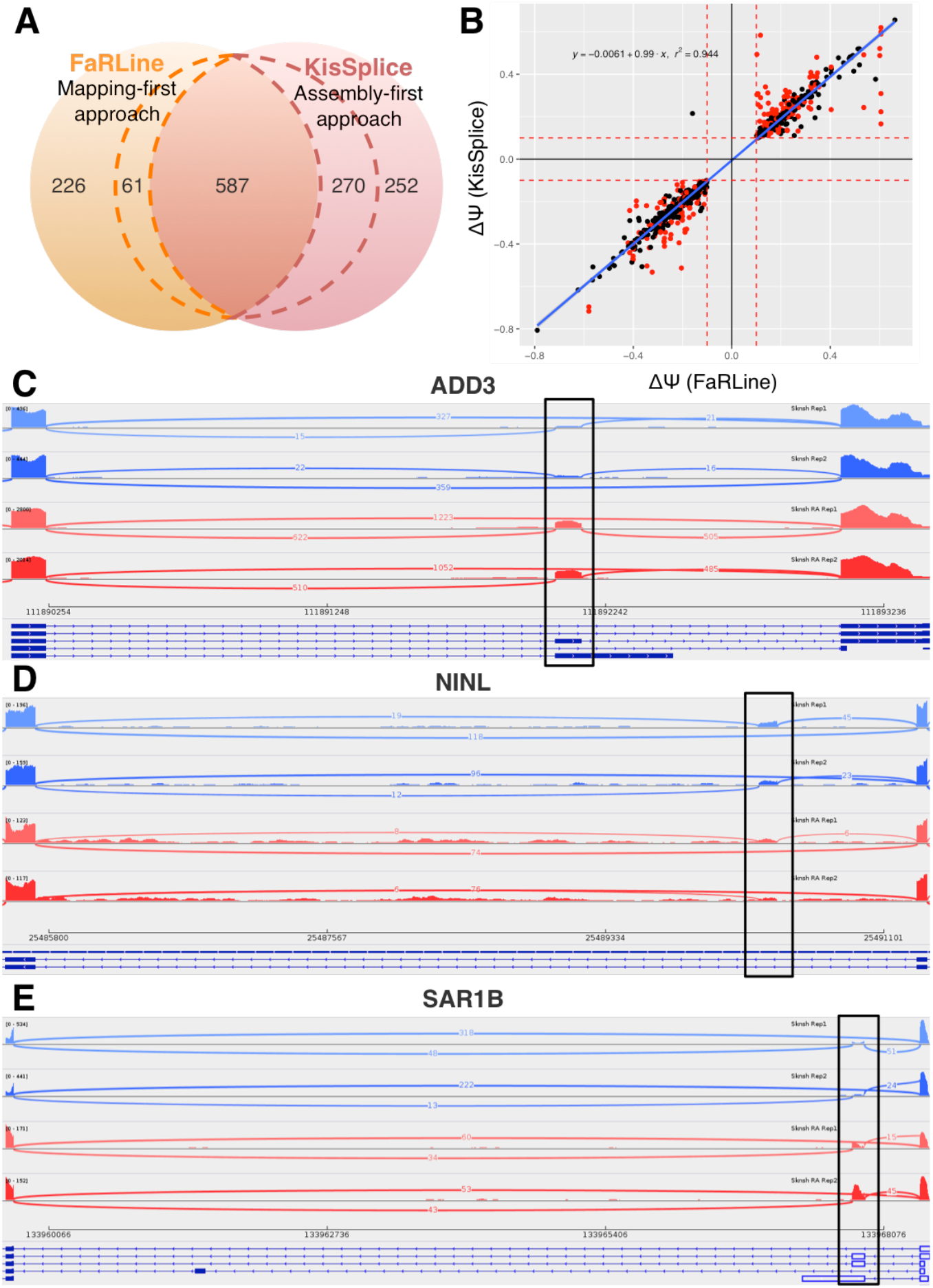
A) Condition-specific variants found by FaRLine, KisSplice or both methods. Within dashed lines are events identified by both approaches but detected as condition-specific by only one approach. B) DeltaPSI as estimated by KisSplice and FaRLine, for events found by both. The red points represent complex events (events for which KisSplice found at least 2 ‘bubbles’). C) Exon 22 of ADD3 is an example of regulated ASE found by both approaches. D) A new exon in intron 5 of NINL gene is found by KisSplice only. The inclusion is differentially regulated between the 2 conditions. E) Because exon 3 of SAR1B is an exonised Alu, only FaRLine finds this ASE. Moreover this exon is significantly more included in the treated cells (SK-N-SH RA).

When compared to the splicing event annotated as reported in Figure 2, we noticed that the proportion of condition-specific events detected by only one method increased, for two main reasons. First, some ASE identified by both approaches were found as differentially included only by one method. This is again due to differences in the quantification of the inclusion levels, especially for ASE with multiple 5’ and 3’ splice sites. Second, some of the exons that were missed out by one method at the identification step happened to be condition specific. This is the case of an exon in NINL intron 5 (Figure 4D), only found by KisSplice because it was not annotated. This is also the case of SAR1B exon 3 (Figure 4E), only found by FaRLine because it overlaps with an Alu element.

The observation that many of the exons detected only by one method are differentially included across conditions confirms that these exons should not be discarded from the analysis. Focusing only on exons predicted by one approach may lead to miss splicing events which are central in the response to treatment.

### 2.5 Overlap with other methods

In a first step, we picked FaRLine and KisSplice as examples of a mapping-first and an assembly-first approach respectively. Clearly, there are other published methods in both categories. MISO is probably the most widely used to annotate AS events. We therefore ran it on the same dataset to check how its predictions overlapped with ours. As shown in Figure 5, 72% of predictions made by MISO were common to both FaRLine and KisSplice, 23% were only common with FaRLine, 2% were only common to KisSplice and the remaining 3% were specific to MISO. Overall, almost all candidates predicted by MISO were also predicted by FaRLine. This large overlap with FaRLine was expected, because both are mapping-first approaches. This also shows that the differences between mapping- and assembly-first approaches reported above are not limited to one mapping-first approach.

**Figure 5:**
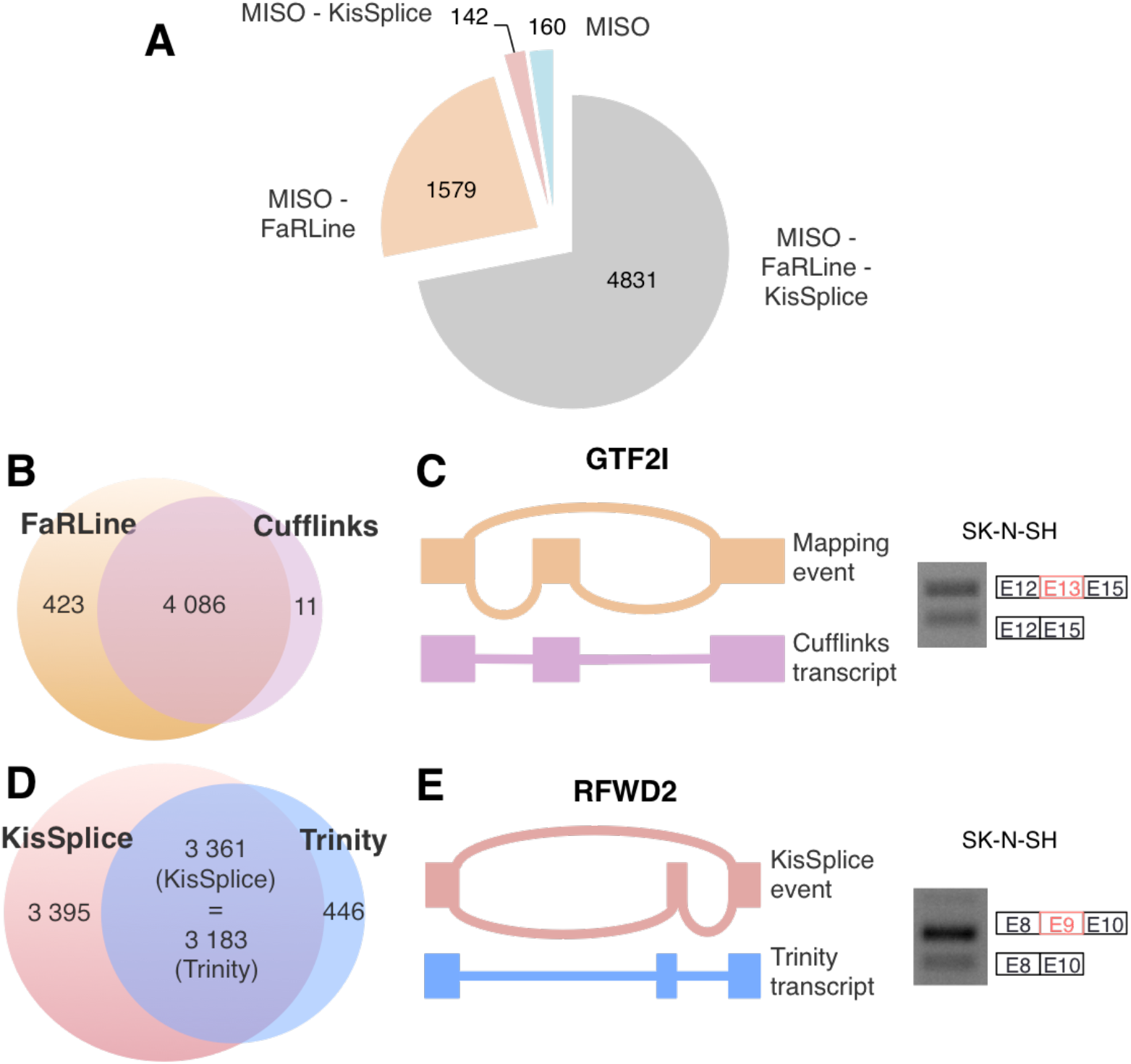
A) 72% of ASE found by MISO are also annotated by FAR-LINE and KisSplice. 23% of MISO’s ASE are also annotated by FAR-LINE while only 3% of MISO’s ASE are also annotated by KisSplice. Finally, only 2% of these ASEs are only annotated by MISO. B) Most of the events annotated by Cufflinks are found by FaRLine. C) GTF2I exon 13 is an example of an ASE annotated by FaRLine but not by Cufflinks. Indeed, Cufflinks only found the inclusion isoform. D) Most of the events annotated by Trinity are also found by KisSplice. But half of the ASE annotated by KisSplice are not found by the global assembler Trinity. E) KisSplice annotate an ASE in the gene RFWD2, while Trinity only found the inclusion variant. The events in panels C and E have been validated by RT-PCR.

Beside exon-centric approaches, which aim at finding the differentially spliced exons, there is also a number of published methods which are isoform-centric and have the more ambitious goal to reconstruct full-length transcripts.

The most widely used mapping-first and isoform-centric approach is Cufflinks [Trapnell et al., 2012] that we compared to FaRLine using the same dataset. As shown in Figure 5D, we found that the vast majority of ASE were predicted by both approaches.

Finally, we compared KisSplice to one of the most widely used de-novo transcriptome assembler, Trinity[Grabherr et al., 2011]. As shown in Figure 5B, most ASE found by Trinity were also found by KisSplice. However, KisSplice was significantly more sensitive. The goal of Trinity is to assemble the major isoforms for each gene, it therefore largely under-estimates alternative splicing, especially inclusion/exclusion of short sequences.

### 2.6 Discussion

*De novo* assembly is usually applied to non-model species where no (good) reference genome is available. We show here that its usage, even when the annotated reference genome is available, offers a number of advantages. We name this approach “assembly-first” because it does use a reference genome, but as late as possible in the process, in order to minimize the *a priori* about which exons should be found.

Using this strategy, we discovered many novel alternatively skipped exons, which were not found by traditional read mapping approaches (Figure 3). While it is believed that the human genome is fully annotated, it is important to underline that we have not yet established a final map of the parts of the genome that can be expressed. It can be anticipated that sequencing of single-cells from different parts of the body will lead to the discovery of a huge diversity and that a substantial number of new exons will be discovered. An example is the case of unannotated skipped exons which overlaps repeat elements. We cannot exclude that this category is currently largely under-annotated.

We also showed that assembly-first approach has the ability to detect splicing variants from paralogous genes (Figure 3). This is because mapping approaches discard reads mapping to multiple genomic locations. Identification of such splicing variants produced from different genomic regions sharing sequence similarities (e.g. paralog genes, pseudogenes) is however very important, since splicing variants generated from paralogous genes but also from pseudogenes may have different biological functions [Poursani et al., 2016].

Conversely, we showed that some ASE were detected only by the mapping-first approach. As shown in Figure 2, we observed that the mapping-first approach has a better ability to detect lowly-expressed splicing variants. Although such lowly-expressed splicing variants are often considered as “noise” or biologically non relevant, caution must be taken with such assumptions for several reasons. First, mRNA expression level is not necessarily correlated with protein expression level. Second, as observed from single-cell transcriptome analyses, some mRNAs can be expressed in few cells, within a cell population (e.g. they are expressed at a specific cell cycle step) and may therefore appear to be expressed at a low level in total RNAs extracted from a mixed cell population [Bacher and Kendziorski, 2016]. Therefore, computational analysis should not systematically discard lowly-expressed splicing variants and filtering these events should depend on the biological questions to address.

We also observed that the mapping-first approach better detects exons corresponding to annotated-repeat elements (Figure 3). While it has been assumed for a long time that repeat elements are “junk”, increasing evidences support important biological functions for such elements. For example, repeat elements like Alu can evolve as exons and the presence of Alu exons in transcripts has been shown to play important regulatory functions [Sorek et al., 2004, Shen et al., 2011].

When two methods have non-overlapping predictions, the temptation could be to focus on exons found by both approaches and discard the others. We argue that this would be a mistake, because these cases can be validated experimentally, and many of them correspond to regulated events, where the inclusion isoform is significantly up or down regulated in presence of a treatment.

In conclusion, combining mapping- and assembly-first approaches allows to detect a larger diversity of splicing variants. This is very important towards the in depth characterization of cellular transcriptome although other approaches are further required to analyze their biological functions.

From a computational perspective, a number of challenges are still ahead of us. The co-development of two approaches enabled us to narrow down the list of difficult instances not properly dealt with by at least one approach, but we cannot exclude that some categories are still missed by both approaches. The categories of challenging cases that we defined in Figure 3: lowly-expressed variants, exonised Alu, complex splicing variants, paralogs have been overlooked up to now. Possibly because they are much harder to detect, they had been assumed to play a minor role in transcriptomes. A number of recent work however argues in the opposite direction.

For exonised ALUs, paralog genes and genes with complex splicing, the possibility to sequence longer reads with third generation techniques [Tilgner et al., 2014, Bolisetty et al., 2015] should prove very helpful. The number of reads obtained with these techniques is however currently much lower than with Illumina, thereby preventing their widespread use for differential splicing, for which the sequencing depth, and not so much the length of the reads, is the critical parameter which conditions the statistical power of the tests. In the coming years, methods combining second and third generation sequencing should enable to obtain significant advances in splicing.

## 3 Material and Methods

### 3.1 FaRLINE and KisSplice

Figure 1 shows the two pipelines that we are comparing. While STAR and TopHat are third-party softwares, we developed the other methods ourselves. KisSplice was introduced in [Sacomoto et al., 2012], kissDE was introduced in [Lopez-Maestre et al., 2016]. KisSplice2RefGenome and FaRLine are methods we introduce in this paper.

For the sake of self-containment, we explain all methods here.

#### 3.1.1 KisSplice

KisSplice is a local transcriptome assembler. As most short reads transcriptome assemblers [Grabherr et al., 2011, Schulz et al., 2012, Robertson et al., 2010], it relies on a De Bruijn graph (DBG). Its originality lies in the fact that it does not try to assemble full-length transcripts. Instead, it assembles the parts of the transcripts where there is a variation in the exon content. By aiming at a simpler goal, it can afford to be more exhaustive and identify more splicing events. The key concept on which KisSplice is built is that variations in the nucleotide content of the transcripts will correspond to specific patterns in the DBG called bubbles. KisSplice’s main algorithmic step therefore consists in enumerating all the bubbles in the graph built from the reads. The sequences corresponding to the two paths of each bubble are then aligned to the reference genome using STAR, and the result of the alignment is analysed using KisSplice2RefGenome to annotate the event.

#### 3.1.2 Alternative splicing events are bubbles in the DBG

Supplementary figure S6 gives a schematic example of two alternative transcripts which differ by the inclusion of one exon. For the sake of simplicity, the example is given for words of length 3, but the reasoning holds for any word length. Each distinct word of length *k* is called a *k*-mer and corresponds to a node of the DBG. There is a directed edge from a node *u* to a node *v* if the last *k* – 1 nucleotides of u are identical to the first *k* – 1 nucleotides of *v*. Each transcript will therefore correspond to a path in the DBG. A pair of internally node-disjoint paths with a common source and target is called a bubble. The smaller path of the bubble corresponds to the exclusion isoform and is composed of all *k*-mers which overlap the junction between the exons flanking the skipped exon. It is therefore usually composed of *k* – 1 *k*-mers. In the special case where the skipped exon shares a prefix with its 3’ flanking exon, or a suffix with its 5’ flanking exon, then the lower path is composed of less than *k* –1 *k*-mers and the *k*-mer which is the source (resp. target) does not correspond anymore to an exonic *k*-mer, but to a junction *k*-mer.

In practice, the DBG is built from the reads, not from the transcripts. The reads stem from possibly all genes expressed in the studied conditions.

Two difficulties arise: reads contain sequencing errors, and repeats may be shared across genes.

#### 3.1.3 Dealing with sequencing errors

As originally described in [Pevzner et al., 2004] and later in [Zerbino and Birney, 2008], sequencing errors generate recognisable structures in De Bruijn graphs, which can be identified and removed. Their systematic removal however prevents assemblers from studying SNPs. A compromise consists in discarding rare *k*-mers from the graph. This is the strategy we use in KisSplice, where we remove all *k*-mers seen only once. This idea is however not sufficient in the context of transcriptome assembly, where the coverage is very uneven and mostly reflects expression levels. For highly expressed genes, several reads may have errors at the same site, generating *k*-mers with a coverage larger than an absolute threshold. We therefore also use a relative cut-off, which we set to 2%. These cut-offs we introduce to remove sequencing errors have an impact on the running time and on the sensitivity. Decreasing them allows to discover rarer isoforms, at the expense of a longer running time.

#### 3.1.4 Dealing with repeats

Repeats are notoriously difficult to assemble in DNAseq data, and were initially thought to be much less problematic in RNAseq, since they are mostly located in introns and intergenic regions. In practice, mRNA extraction protocols are not perfect, and a fraction of pre-mRNA remains (typically 5% for total polyA+ RNA [Tilgner et al., 2012]). Each intron is covered by few reads, but if a repeat is present in many introns, then this repeat will obtain a high coverage and will correspond to very dense subgraphs in the De Bruijn graph built from the reads. The traversal of such subgraphs to enumerate all the bubbles they contain is long and mostly fruitless. We showed in [Sacomoto et al., 2014] that an effective strategy to deal with this issue is to enumerate only bubbles which have at most *b* branches. In practice, we set *b* to 5. Increasing *b* will increase the running time, but allow to find more repeat-associated alternative splicing events. Bubbles which do not correspond to true AS events can be filtered out at the mapping step.

#### 3.1.5 Annotating the events with KisSplice2RefGenome

Bubbles found by KisSplice are mapped to the reference genome using STAR, with its default settings, which means that in case of multi-mappings, STAR reports all equally best matches. The mapping results are then analysed by KisSplice2RefGenome. At this stage, bubbles are classified in sub-types depending on the number of blocks obtained when mapping each path of the bubble to the genome (Supplementary Figure S7). For exon skip-pings, the longer path of the bubble corresponds to 3 blocks, while the lower path corresponds to 2 blocks. The splice sites are located and compared to the annotations. Events with novel splice sites are reported explicitly in the output of the program.

In the case where the bubble corresponds to a genomic insertion or deletion, it exhibits a specific pattern in terms of block numbers and is reported separately.

In the case where the bubble maps to two locations on the genome, a distinction is made between the case of exact repeats where both paths map to both locations and inexact repeats where each path maps to a distinct location (Supplementary Figure S8). The cases of exact repeats corresponds to recent paralogs.

#### 3.1.6 FaRLine

##### FasterDB EnsEMBL r75 annotation

FasterDB RNAseq Pipeline, FaRLine, use the FasterDB-based EnsEMBL r75 annotation database. FasterDB is a database containing all annotated human splicing variants [Mallinjoud et al., 2014].

The genomic exons are defined by projecting the transcript exons (Supplementary Figure S9). First, the transcript exons are grouped by position. Then each group of exons define a projected exon with the following rules:

- The start is the smallest start of the non-first-exon of the group.
- The end is the highest end of the non-last-exon of the group that ends before the start of the next group of exons.

When the most frequent event annotated in the transcrits is an intron retention, the projected genomic exon is defined as a combination of the two exons the intron retained. In supplementary figure S9, the exons 5 and 6 and the intron 5 are considered as one unique exon. As events included in an exon are overseen, this results in some events being missed.

##### Mapping

The first step of FaRLine is to map the reads to a reference genome. This step is done using Tophat-2.0.11 [Trapnell et al., 2012]. `tophat --min-intron-length 30 --max-intron-length 1200000 \ -p 8 [--solexa1.3-quals for Sknsh_rep1 and Sknsh_rep2] \ --transcriptome-index`

A transcriptome index has been built by TopHat using EnsEMBL r75 annotations in gtf format. When a transcriptome index is used, the mapping steps are modified: instead of aligning first to the genome, which is the default behavior, TopHat uses Bowtie to align the reads to the transcript sequences first, then align the remaining unmapped reads to the genome. Minimal and maximal intron lengths have been modified (default 70 and 500000) to maximize the number of junctions detected, according to the statistics provided by FasterDB EnsEMBL r75 annotations.

The resulting alignment files have been filtered using samtools 0.1.19 [Li et al., 2009].

`samtools view -F 260 -f 1 -q 10 -b`

With this step, only the primary alignments are kept. The minimum read alignment quality was set up so that multi-mapping reads were removed from the alignment file.

##### Annotation and quantification of alternative splicing events

We wrote custom perl scripts, based on the FasterDB-based EnsEMBL r75 annotation database. For each gene, all the reads with at least one base overlapping the gene from the start to the end coordinates are retrieved. CIGAR strings are then used to retrieve the alignments blocks. Junction reads are identified by the presence of at least one ’N’ letter in the CIGAR. Junction reads were filtered if:

- More than 10% of soft-clipping was detected in the alignment (it should not be the case with TopHat)
- An indel was close to the junction site, as it would make the junction position uncertain

Junction read alignments are then processed block by block sequentially from left to right. Alignment blocks under 4bp on read extremities are removed from the reads as we considered it is not sufficient to identify correctly the mapping localization. Then each block is compared to FasterDB annotations to check if the block boundaries correspond to known exons annotated in FasterDB, or to a putative new acceptor or donor site. First and last alignment blocks for each read must overlap one and only one exon for a read to be considered. For the inner blocks, if alignment blocks map to a succession of exons and introns, it is considered as an intron retention. However, as the read size is only 76bp, this should not happen often. For the acceptors and donors, we also added a supplementary filter. If a new donor is identified within a junction, we check if the junction also has an acceptor identified of the same length +/−1bp on the other side of the junction, showing most probably a problem of mapping. Once all the blocks are identified, the block annotations are used to annotate putative alternative splicing events: alternative skipped exon, multiple exon skipping, acceptor, or donor sites.

Once all the junction reads are processed, the alternative splicing events identified are pooled and the read participating to each event are quantified, as well as the known exon-exon junction. If an exon-exon junction is annotated with multiple known acceptors and/or donors, all the possible junction reads are quantified and summed up. To fasten the quantification step, a junction coordinate file with the corresponding read numbers is produced from the read alignment using the same filters than described above and will be used for all the quantification tools: junction, exon skipping, acceptor and donor.

A challenge in defining the alternative skipped exon events is to identify the flanking exons. In the first version of FaRLine, these flankings exons were defined as the closest annotated genomic exons. This rule led to miss a lot of ASE events. So to define the flanking exons, we use the information contained in the transcripts and in the reads. We list each junction skipping an exon covered by at least one read. If this junction is annotated in the transcripts, we extract all annotated events containing this junction. Else, we annotate the event with the longest covered inclusion isoform. It allows FaRLine to be more robust to the incompleteness of the annotation compared to other methods, like MISO. Panel B of supplementary figure S2 gives an example of an ASE reported by FaRLine but not by MISO because the inclusion isoform is not annotated in the transcripts.

##### Comparison with STAR

We also mapped the reads with STAR, ran FaRLine on this alignments and compared the predicted skipped exon with KisSplice. The main results are similar to what we found with TopHat. Indeed, without any filter, 69% of ASE annotated by KisSplice are also found by FaRLine and 24% of FaRLine’s event by KisSplice (compared to 68% and 24% respectively for the mapping with TopHat). When we filter out the events with an unfrequent variant, we show that approximately 70% of predicted ASE are found by both approaches.

#### 3.1.7 Differential analysis

Both pipelines perform ASE detection and quantification. The last step of the pipelines is the differential analysis of the expression levels of the variants. This task is performed using the kissDE [Lopez-Maestre et al., 2016] R package, which takes as input a table of read counts as in Figure S10, and outputs a p-value and a DeltaPSI (Percent Spliced In).

Our statistical analysis adopted the framework of count regression with Negative Binomial distribution. We considered a 2-way design with interaction, with *isoforms* and *experimental conditions* as main effects. Following the Generalized Linear Model framework, the expected intensity of the signal was denoted by *λ*_*ijk*_ and was decomposed as:

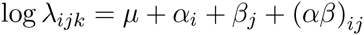

where *μ* is the local mean expression of the gene, *α*_*i*_ the contribution of splicing variant *i* on the expression, *β*_*j*_ the contribution of condition *j* to the total expression, and (*αβ*)_*ij*_ the interaction term. The target hypothesis was *H*_0_: {(*αβ*)_*ij*_ = 0} *i.e.* no interaction between the variant and the condition. If this interaction term is not null, a differential usage of a variant across conditions occurred. The test was performed using a Likelihood Ratio Test with one degree of freedom. To account for multiple testing, p-values were adjusted with a 5% false discovery rate (FDR) following a Benjamini-Hochberg procedure [Benjamini and Hochberg, 1995].

In addition to adjusted p-values, we report a measure of the magnitude of the effect. The measure we provide is based on the Percent Spliced In (PSI) calculated for a pair of variants:

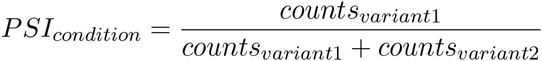

If counts for a variant are below a threshold, then the PSI is not calculated. This prevents from over-interpreting large magnitudes derived from low counts. When several replicates are available for a condition, then a PSI is computed for each replicate, and then we calculate their mean.

Finally, we output the DeltaPSI:

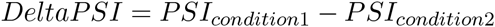

unless one of the mean PSI of a condition could not be estimated. The higher the DeltaPSI, the stronger the effect. In practice, we consider only DeltaPSI larger than 0.1, a threshold below which it is extremely difficult to perform any experimental validation.

### 3.2 SKNSH dataset

We downloaded a total of 959M reads from http://genome.crg.es/encode_RNA_dashboard/hg19/. They correspond to long polyA+ RNAs generated by the Gingeras lab, and are also accessible with the following accession numbers (ENCSR000CPN – SRA: SRR315315, SRR315316 and ENCSR000CTT -SRA: SRR534309, SRR534310). For cell lines treated by retinoic acid, the reads were 76nt long, while they were 100nt long for the non treated cells. Hence we trimmed all reads to 76nt.

### 3.3 Computational requirements

FaRLine took 45 hours and 10 Go of RAM. The time-limiting step was TopHat2, which took 41 hours, even parallelised on 8 cores. When STAR was tested instead of TopHat2, it took 4 hours, but 30 Go of RAM. KisSplice took 30 hours and 10Go RAM. The RAM-limiting step was STAR which took 30Go of RAM. All the steps of the pipelines can be reproduced using the following tutorial: http://kissplice.prabi.fr/sknsh/.

### 3.4 Experimental Validation

SK-N-SH cells were purchased from the American Type Culture Collection (ATCC) and cultured using EMEM medium (ATCC) complemented with 10% FBS (Thermo Fisher Scientific). Cells were differentiated for 48h using 6*µ*M of all-trans retinoic acid (Sigma-Aldrich).

After harvesting, total RNA were extracted using Tripure isolation reagent (Sigma-Aldrich), treated with DNase I (DNAfree, Ambion) for 30 min at 37°C and reverse-transcribed (RT) using M-MLV reverse transcriptase and random primers (Invitrogen). Before PCR, all RT reaction mixtures were diluted at 2.5 ng*µ*L of initial RNA. PCR reactions were performed using GoTaq polymerase (Promega).

## 4 Acknowledgments

This work was funded by the ANR-12-BS02-0008 (Colib’read) by the ABS4NGS ANR project (ANR-11-BINF-0001-06), Action n3.6 Plan Cancer 2009–2013, Fondation ARC (Programme Labellisé Fondation ARC 2014, PGA120140200853) and INCa (2014-154). Doctoral fellowships from ARC 1 – Région Rhône-Alpes (C.B.P), Science Without Borders – CNPq – Brazil (L.L. – grant process number 203362/2014-4), ARS Rhône-Alpes (A.R.) and post-doctoral fellowships from Fondation ARC (M.P.L).

This work was performed on the computing facilities of the computing center LBBE/PRABI and the PSMN (Pole Scientifique de Modelisation Numerique) computing center of ENS de Lyon.

---

**Figure S1:**
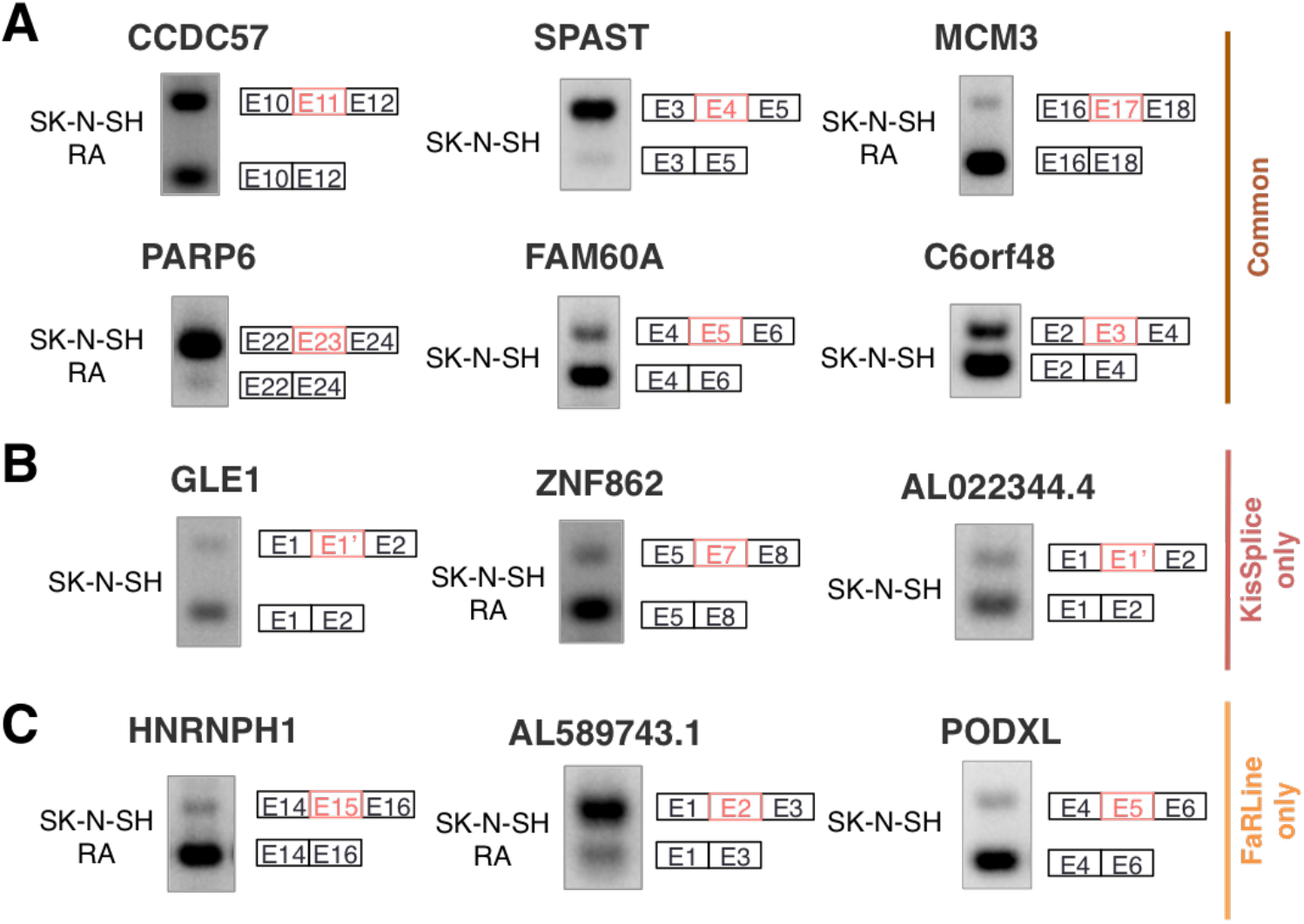
rt-PCR validations of events found by both approaches (A), only by KisSplice (B) and only by FaRLine (C).

**Figure S2:**
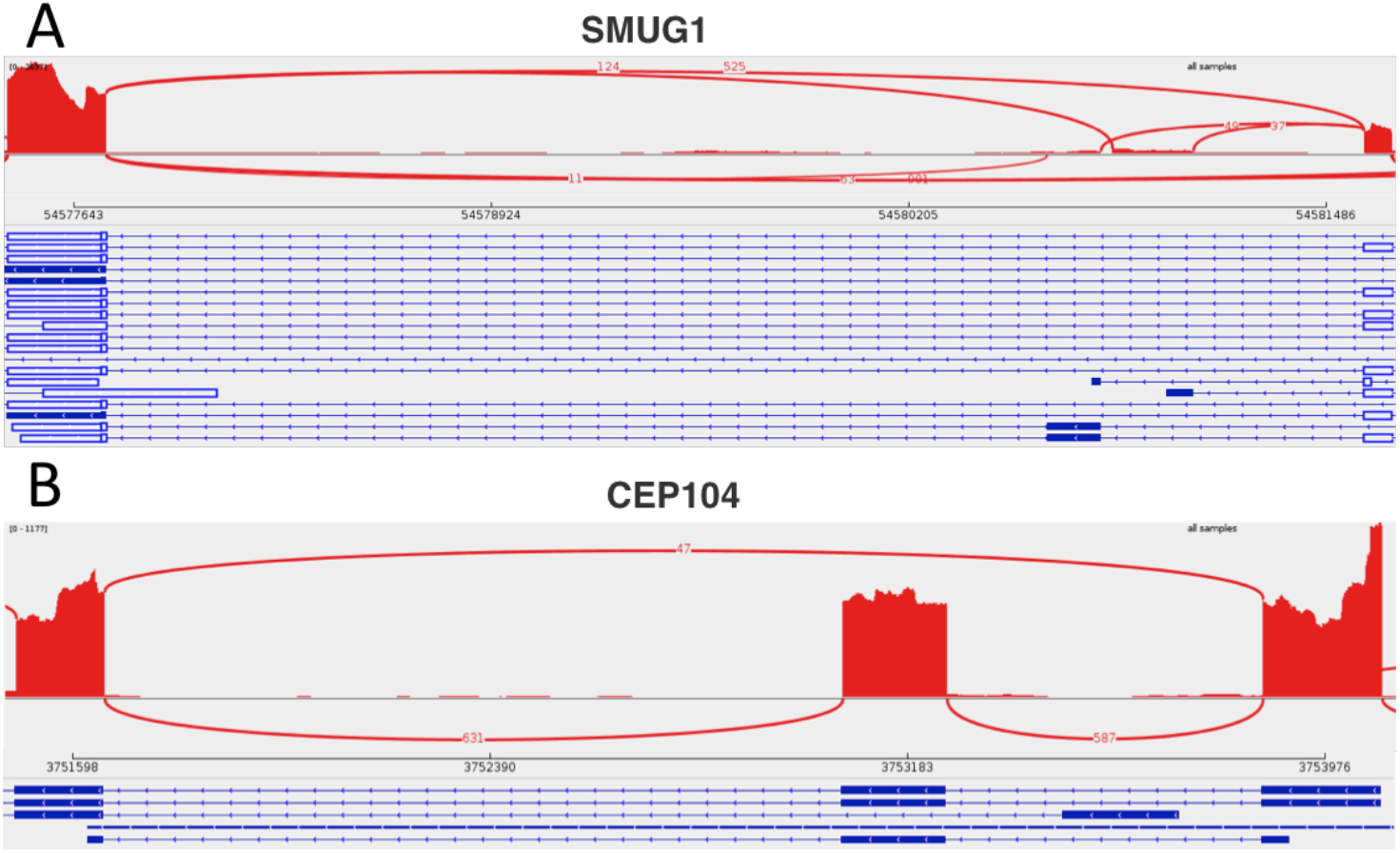
Examples of exon skipping inside a complex event. A) The exon 5 of SMUG1 gene is reported as skipped by KisSplice with exons 4 and 7 as flanking exons. This event is not found by FaRLine because the inclusion isoform is not annotated in the transcrits. B) Exon 12 of CEP104 gene is reported as skipped by FaRLine even if the exclusion isoform is not present in the annotation. However, MISO does not find this exon skipping.

**Figure S3:**
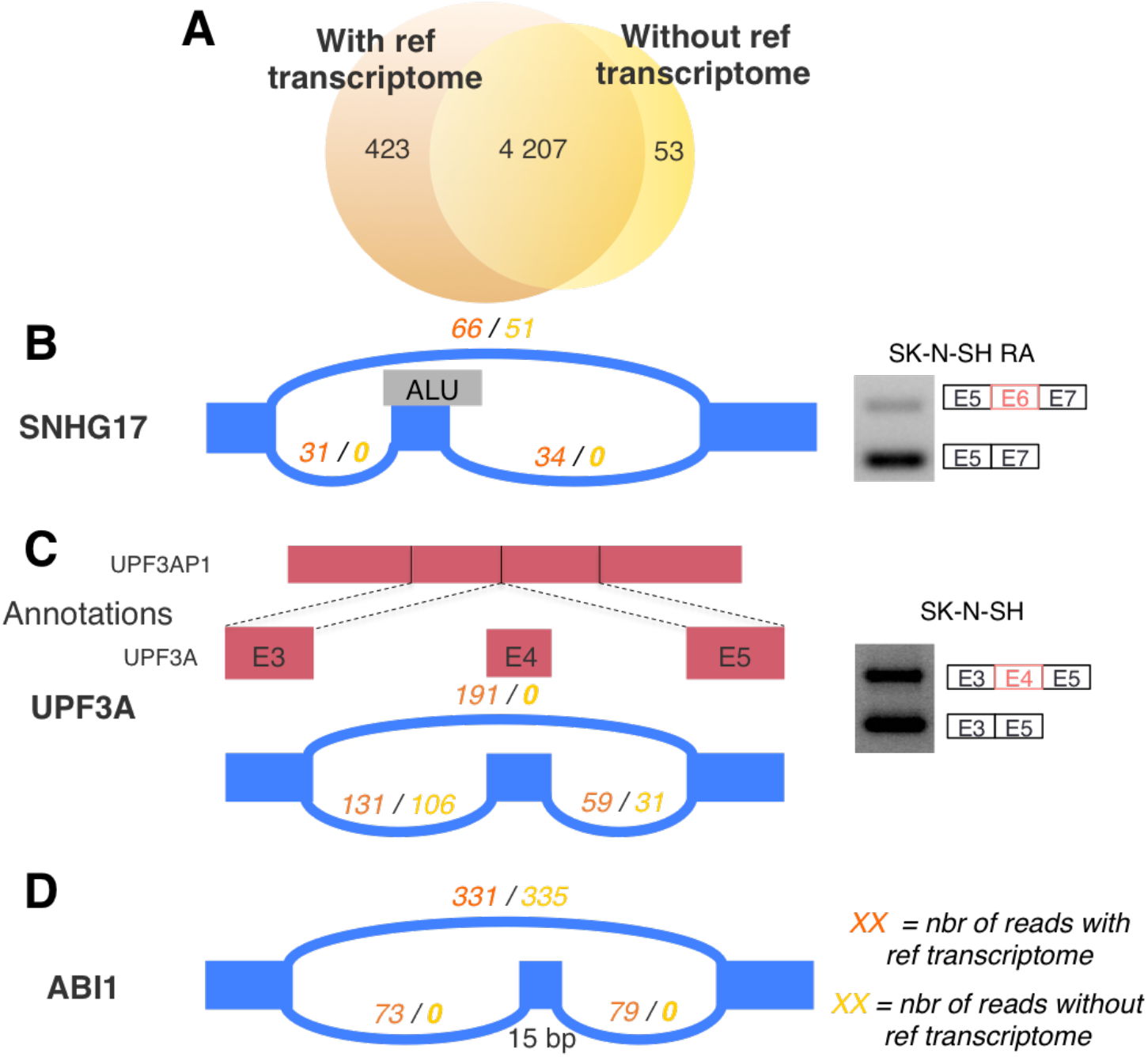
Comparison of the mapping-first approach FaRLine with or without an annotation provided to the mapper (i.e. with/without reference transcriptome). A) More ASE are annotated when an annotation available. Panels B to D show examples of events only found by the mapping-first method when an annotation is provided to the mapper. B) The first category, represented by the gene SNHG17, includes exons containing repeats like ALU elements. C) Genes with a retrotransposed pseudogene, as UPF3A, represent the second category and are more difficult to find when no annotation is available. D) Short exons (less than 20bp), like exon 5 of the gene ABI1, compose the third category.

**Figure S4:**
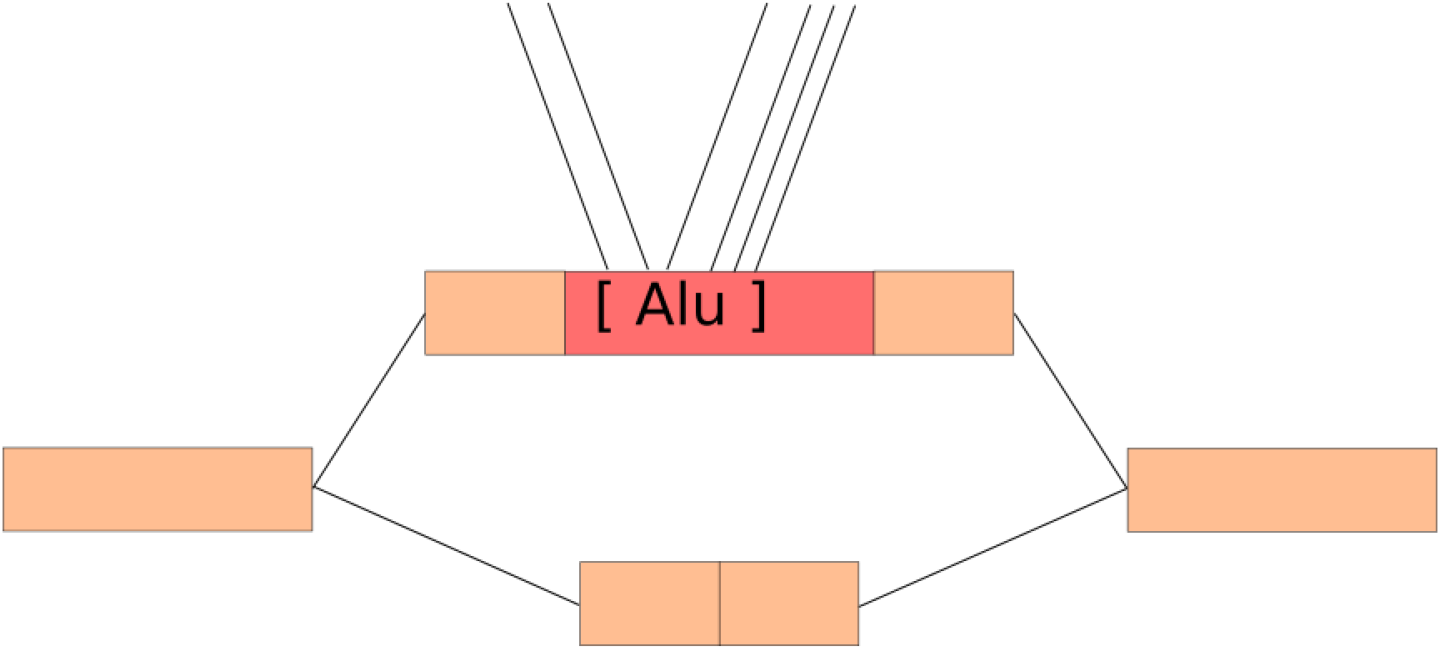
Example of a bubble containing an Alu. Repeated events such as Alu are expected to be present in several copies in the reads. Thus, when the graph is constructed, edges link different copies of Alu. Because a bubble with more than 5 edges within one of its paths is not enumerated by KisSplice, this case is not annotated by the assembly-first approach.

**Figure S5:**
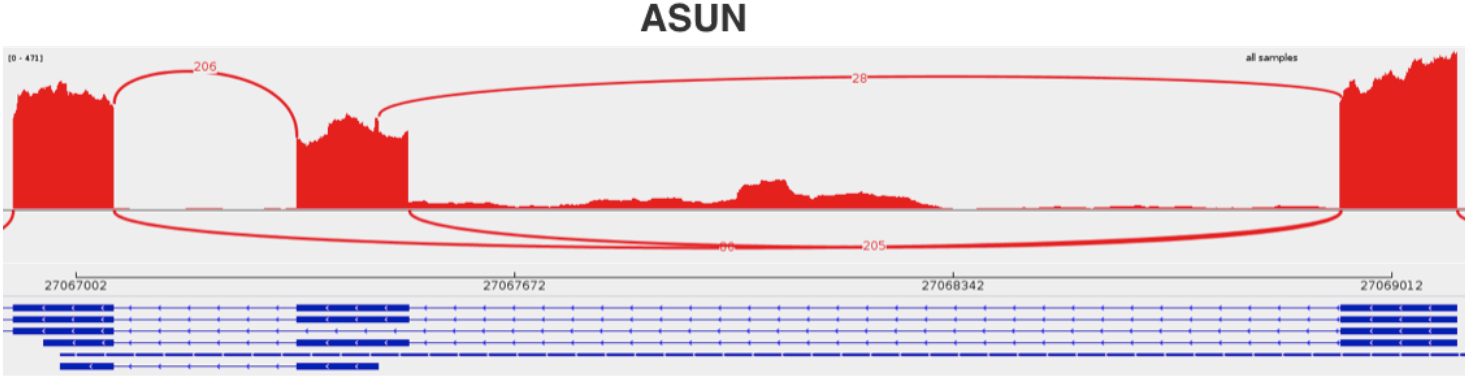
Example of an exon skipping with two alternative donor sites. It is reported as one event by FaRLine and two events by KisSplice.

**Figure S6:**
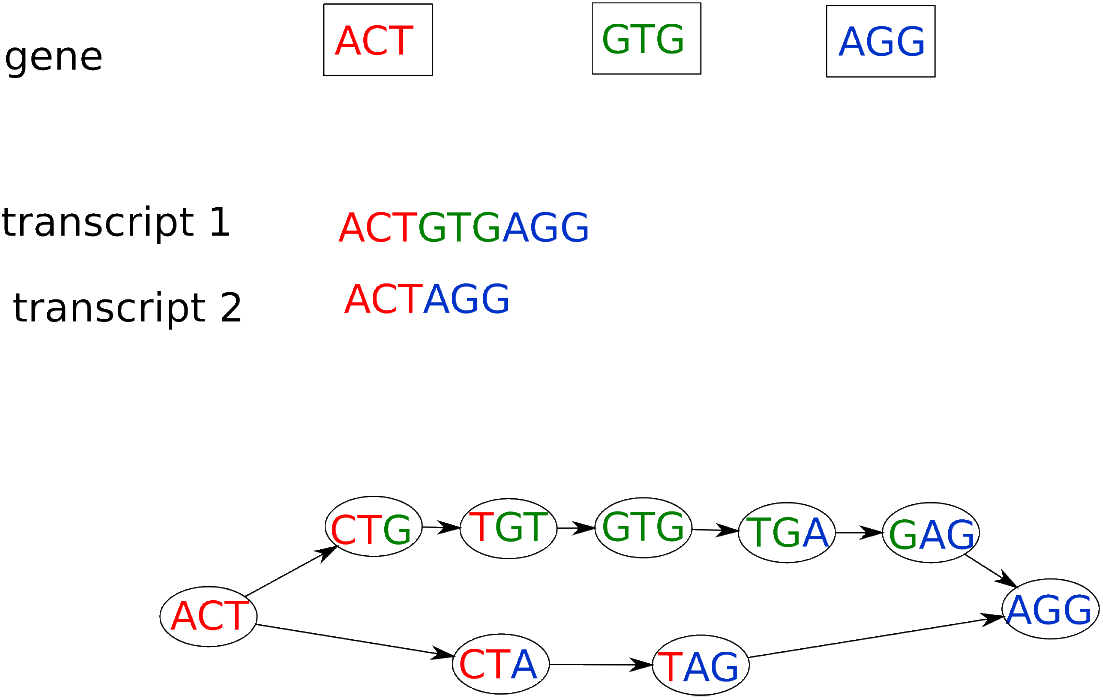
A schematic gene with three exons producing two alternative transcripts. The De Bruijn graph built from the sequences of the transcripts corresponds to a bubble. The upper path spells the skipped exon and its flanking junctions while the lower path spells the junction of the exclusion isoform and has a predictable length.

**Figure S7:**
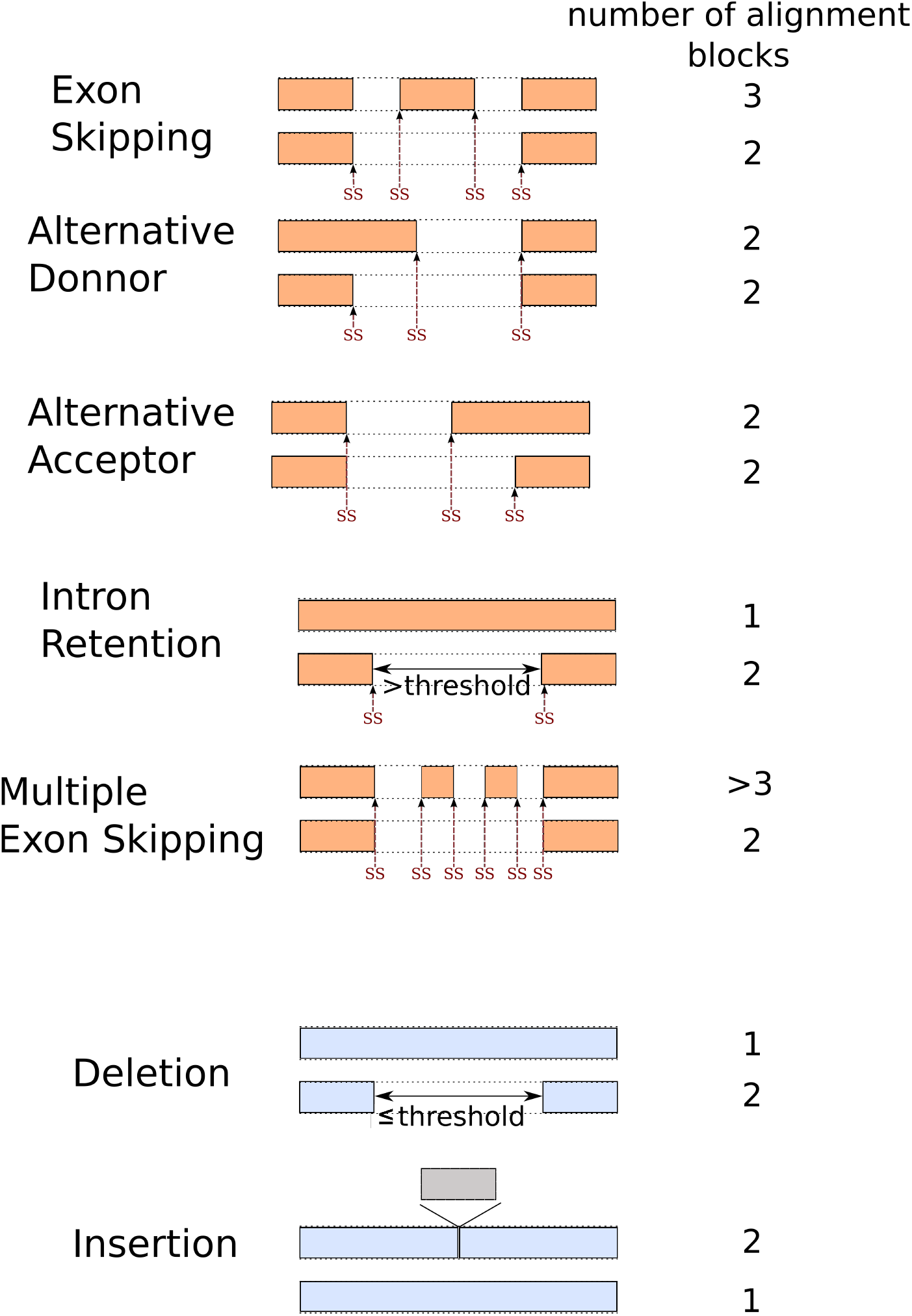
Classification of KisSplice events according to the number of blocks in which they map to the reference genome. Paths representing variants of an event are mapped on the reference. Spliced mapping results in blocks, events are then classified by KisSplice2RefGenome according to the block mapping patterns. (Putative) splice sites are noted by SS in red. In addition to alternative splicing, some indels are filtered through this step as they correspond to specific block patterns too.

**Figure S8:**
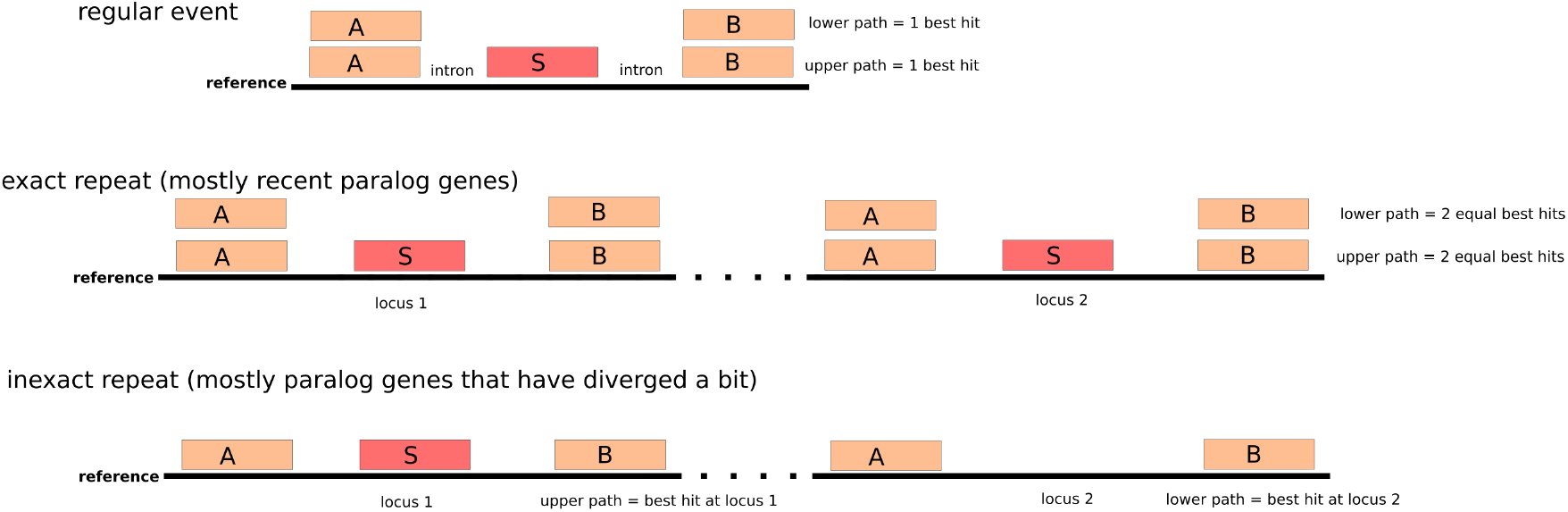
Dealing with repeats in KisSplice2RefGenome. If the two variants (i.e. paths) both map on different copies (exact repeat), we classify it as a recent paralog. On the contrary if each variant maps on a different locus, we consider the event as coming from an inexact repeat. This category represents mostly paralogs that have diverged.

**Figure S9:**
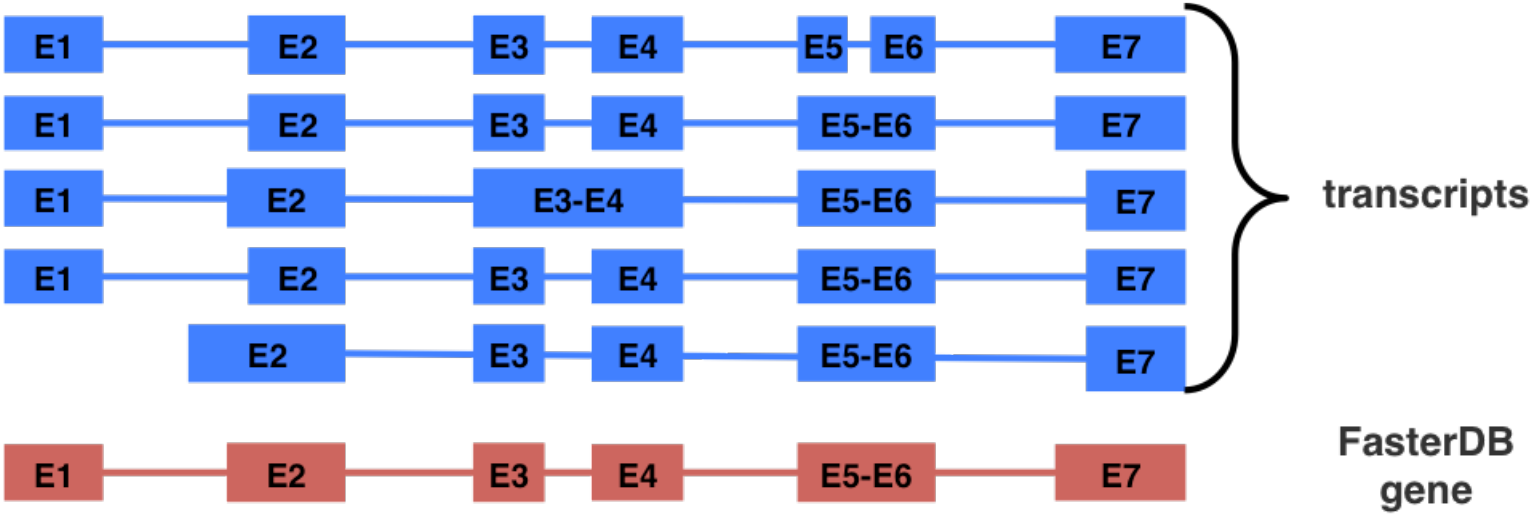
FasterDB exons are defined as the projection of the longer or most frequent exon in the transcripts (except for alternative first or last exons). The whole analysis done with FaRLine is based on these exons.

**Figure S10:**
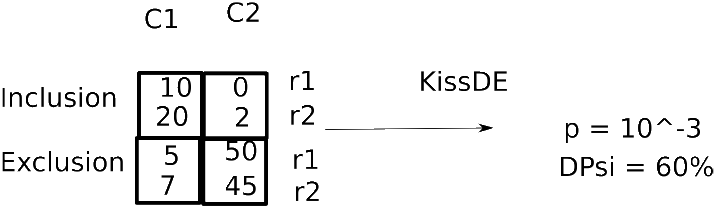
Input and output of the differential analysis. Counts for each replicate of each condition are computed by FaRLine or KisSplice. These counts together with the experimental plan are the input of kissDE. In the example, we show counts for one single event, in practice kissDE tests all events discovered by one method to spot the differential splicing events. Provided at least two replicates are available per condition, kissDE computes p-values and DeltaPSI per event, and results are ranked using these two metrics.

